# Key common genes and pathways in ulcerative colitis and ankylosing spondylitis based on bioinformatics analysis

**DOI:** 10.1101/2023.04.20.537616

**Authors:** Lin Li, Fuzhen Li, Kunpeng Xie, Pengyi Zhou, Haiyan Zhu, Liping Du, Peizeng Yang, Xuemin Jin

**Affiliations:** Department of Ophthalmology, The First Affiliated Hospital of Zhengzhou University, Henan Province Eye Hospital, Henan International Joint Research Laboratory for Ocular Immunology and Retinal Injury Repair, Zhengzhou, Henan, P.R. China; The First Affiliated Hospital of Chongqing Medical University, Chongqing Key Laboratory of Ophthalmology and Chongqing Eye Institute, Chongqing, China

## Abstract

Associations between ulcerative colitis (UC) and ankylosing spondylitis (AS) have been observed in multiple studies, but the common etiology of UC and AS remain unknown. Thus, the current research was conducted to investigate the shared genes and relevant mechanisms in UC and AS. GSE87466 and GSE25101 datasets were used to identify DEGs involved in UC and AS, respectively. The clusterProfiler R package was utilized to detect the biological processes of DEGs in UC and AS. The performance of common DEGs in distinguishing UC or AS samples from control ones were evaluated by ROC curves. The miRWalk, Cistrome and TransmiR database were utilized to construct the network of TF-miRNA-diagnostic biomarker. GSEA method and CTD database were used to investigate the common KEGG pathways shared by UC and AS. In addition, CTD database was also used to detect the interaction score between diagnostic biomarkers and diseases associated with UC or AS. Moreover, prospective diagnostic biomarker-targeting drugs were identified using the DGIdb database. A total of 20 common DEGs were obtained by analyzing data in GSE97466 and GSE25101 datasets. ROC curves revealed that GMFG, GNG11, CLEC4D, CMTM2, VAMP5, S100A8, S100A12 and DGKQ may serve as diagnostic biomarkers for the individuals with AS and UC. A network of TF-miRNA-diagnostic biomarker, composed of 212 nodes and 721 edges, was constructed and visualized by Cytoscape software. Toll-like receptor signaling pathway, antigen processing and presentation, allograft rejection, viral myocarditis, pathways in cancer, graft versus host disease and natural killer cell mediated cytotoxicity were identified as common pathways in UC and AS. For the first time, our study identified 8 common key genes and 7 common pathways in UC and AS. These findings may help to clarify the relationship between UC and AS, and provide guidance in the diagnosis and treatment of UC and AS patients.

**Author Summary:** Ankylosing spondylitis (AS) and Ulcerative colitis (UC) are two types of autoimmune diseases that often co-occur. The simultaneous onset of both diseases often results in severe clinical manifestations and limited therapeutic efficacy. So there is an urgent need to gain a deeper understanding of the causes of UC and AS in order to develop more effective treatment strategies. In this study, we explored the genes and pathways commonly involved in two diseases, constructed a transcriptional network, and further investigated potential drugs. The discoveries could potentially offer insights into the connection between UC and AS, and assist in identifying and managing patients with UC and AS.

## Introduction

Ulcerative colitis (UC), classified as an inflammatory bowel disease (IBD), impacts the digestive system and causes persistent inflammation within the inner lining of the large intestine[1]. In recent years, many studies have been conducted to gain further insights and understandings into this debilitating condition. Symptoms of UC vary from person to person, but common signs and symptoms may include diarrhea, rectal bleeding, abdominal cramps and pain, unintended weight loss, and anemia[2]. In addition, some people with UC may also experience joint pain, skin rashes, and eye irritation[3]. Between 20 and 250 instances of UC are believed to occur for every 100,000 people, depending on the population and geographic region. Several studies reveal the prevalence of UC has doubled in the last two decades, indicating that the condition is becoming more common worldwide[3]. Multiple factors, including microbial infection, immune response, endocrine abnormity and biological heritage, have been linked to the development of UC[2]. The treatment of UC primarily concentrates on reducing inflammation, relieving symptoms, and maintaining remission[4]. In order to develop more effective treatment strategies, further understanding the causes and symptoms of UC is still urgent.

Ankylosing spondylitis (AS) is a type of chronic and inflammatory arthritis that mainly targeting the spine and sacroiliac joints[5]. The condition is distinguished by progressive stiffness and fusion of the spine, which causes severe back pain, limited mobility, and disability[6]. Other associated symptoms may include fever, weight loss, joint swelling, and eye inflammation. Studies have shown that the prevalence of AS is 0.1-0.2% in the general population, with a greater frequency in specific populations[7]. AS is a complex disorder that can significantly impact patients’ quality of life, and the causes of this condition remain largely unknown. However, genetic and environmental factors may play a part[7]. Current treatments for the disease are still limited to managing the symptoms, such as pain and stiffness, by taking medications, engaging in physical activity and other lifestyle adjustments[8]. Therefore, further research is required to better understand the pathogenesis of AS and to develop strategies to improve the management of this condition.

Several studies have proved the link between AS and UC. 5-10% of AS patients are also diagnosed with IBD, implying that AS may be an independent risk factor for IBD[9]. Furthermore, UC can often lead to Sacroiliac arthritis which is characterized by lumbosacral pain and limited mobility. The prevalence of Sacroiliac arthritis in individuals with UC is estimated to be around 20%[10]. Symptoms of Sacroiliac arthritis may improve or disappear along with UC. However, when UC is combined with AS, patients usually experience severe joint symptoms, and inadequate treatment may lead to deformity and spinal rigidity[11]. Therefore, early diagnosis is crucial for the prognosis of patients suffering from both disorders. The diagnosis of AS combined with UC is often unclear based on clinical manifestations. Thus, exploring the potential common markers and mechanisms of these two diseases is vital.

The common processes underlying the prevalence of AS and UC are still unknown. Our research aims to investigate the key genes and associated pathways implicated in both disorders. This will assist in setting up the basis for a more comprehensive understanding of the connection between AS and UC.

## Results

### DEGs involved in UC were associated with immune and extracellular matrix

The analysis conducted in GSE87466 revealed a set of 3,123 DEGs (1,749 up-regulated genes and 1,374 down-regulated genes) between controls and UC samples (Fig 1A and S1 Table). The expressions of those DEGs were displayed in the heatmap (Fig 1B). To detect the biological effect of DEGs, GO and KEGG pathway analyses were conducted. A total of 1,463 BPs, 76 CCs, 146 MFs (S2 Table) and 67 KEGG pathways (S3 Table) were significantly enriched. As demonstrated in Fig 1C and 1D, DEGs exhibited significant enrichment in immune and extracellular matrix related biological functions, including extracellular matrix organization and positive regulation of cell adhesion in terms of BPs, external side of plasma membrane and collagen-containing extracellular matrix in terms of CCs, as well as extracellular matrix structural constituent and cytokine activity in terms of MFs. In addition, the DEGs were found to be enriched in KEGG pathways, including cell adhesion molecules and cytokine-cytokine receptor interaction.

**Fig 1.**
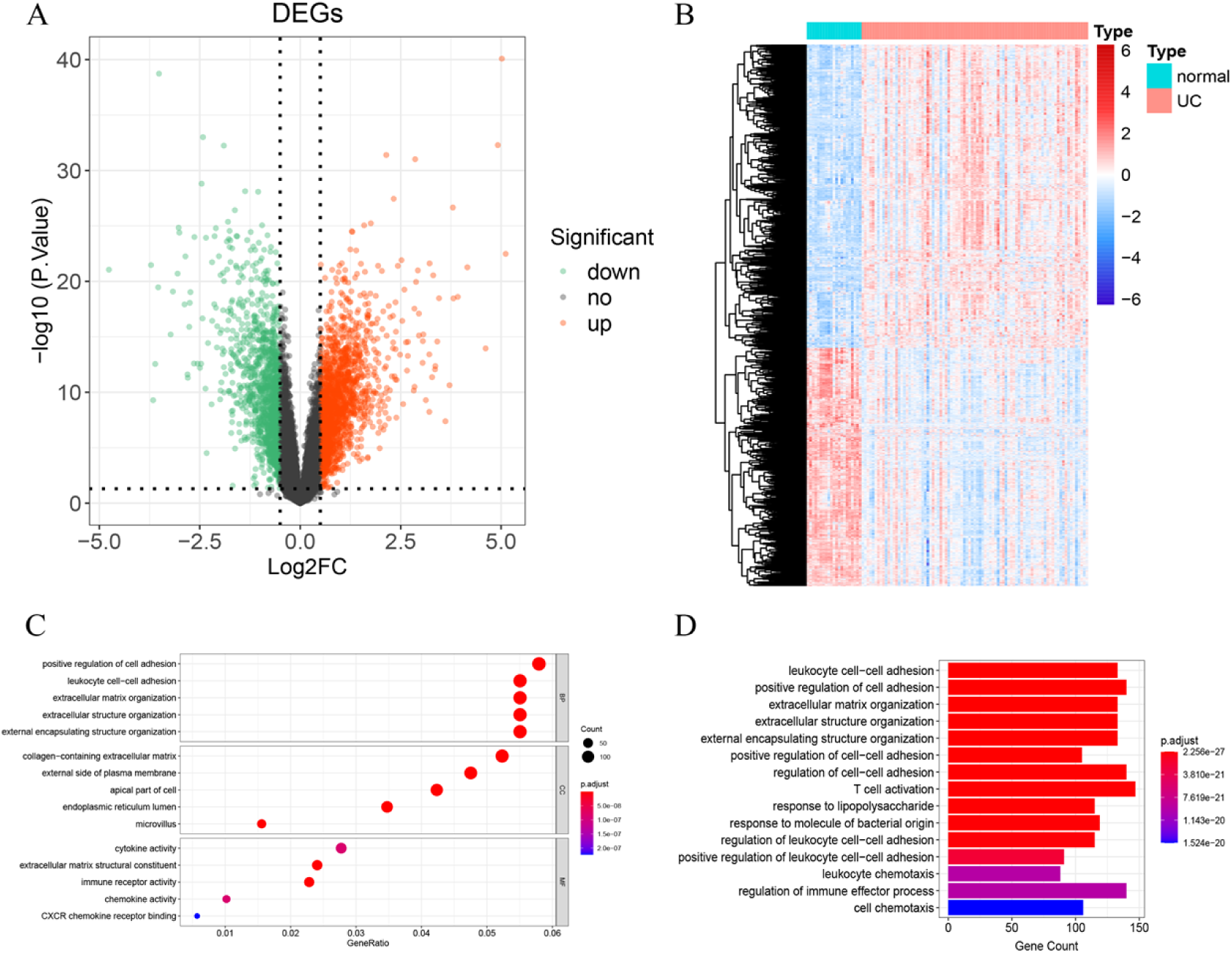
Identification and functional enrichment analysis of DEGs involved in UC. A. The volcano plot of DEGs identified in GSE87466 (n=108, p < 0.05) ; B. The heatmap plot of DEGs identified in GSE87466; C. The volcano plot of DEGs identified in GSE87466 (p < 0.05) ; D. Enrichment analysis of GSE87466.

### DEGs involved in AS were associated with protein biogenesis, oxidative stress and neurodegeneration

The analysis conducted in GSE25101 revealed a set of 137 DEGs (100 up-regulated genes and 37 down-regulated genes) between controls and AS samples (Fig 2A and S4 Table). The expressions of those DEGs were displayed in the the heatmap (Fig 2B). To detect the biological effect of DEGs, GO and KEGG pathway analyses were conducted. A total of 80 BPs, 28 CCs, 18 MFs (S5 Table) and 16 KEGG pathways (S6 Table) were significantly enriched. As demonstrated in Fig 2C and 2D, DEGs exhibited significant enrichment in protein biogenesis, oxidative stress and neurodegeneration related biological functions, including cotranslational protein targeting to membrane and establishment of protein localization to endoplasmic reticulum in terms of BPs, ribosomal subunit and respiratory chain complex in terms of CCs, as well as structural constituent of ribosome and RAGE receptor binding in terms of MFs. In addition, the DEGs were found to be enriched in KEGG pathways, including ribosome, Parkinson disease and oxidative phosphorylation.

**Fig 2.**
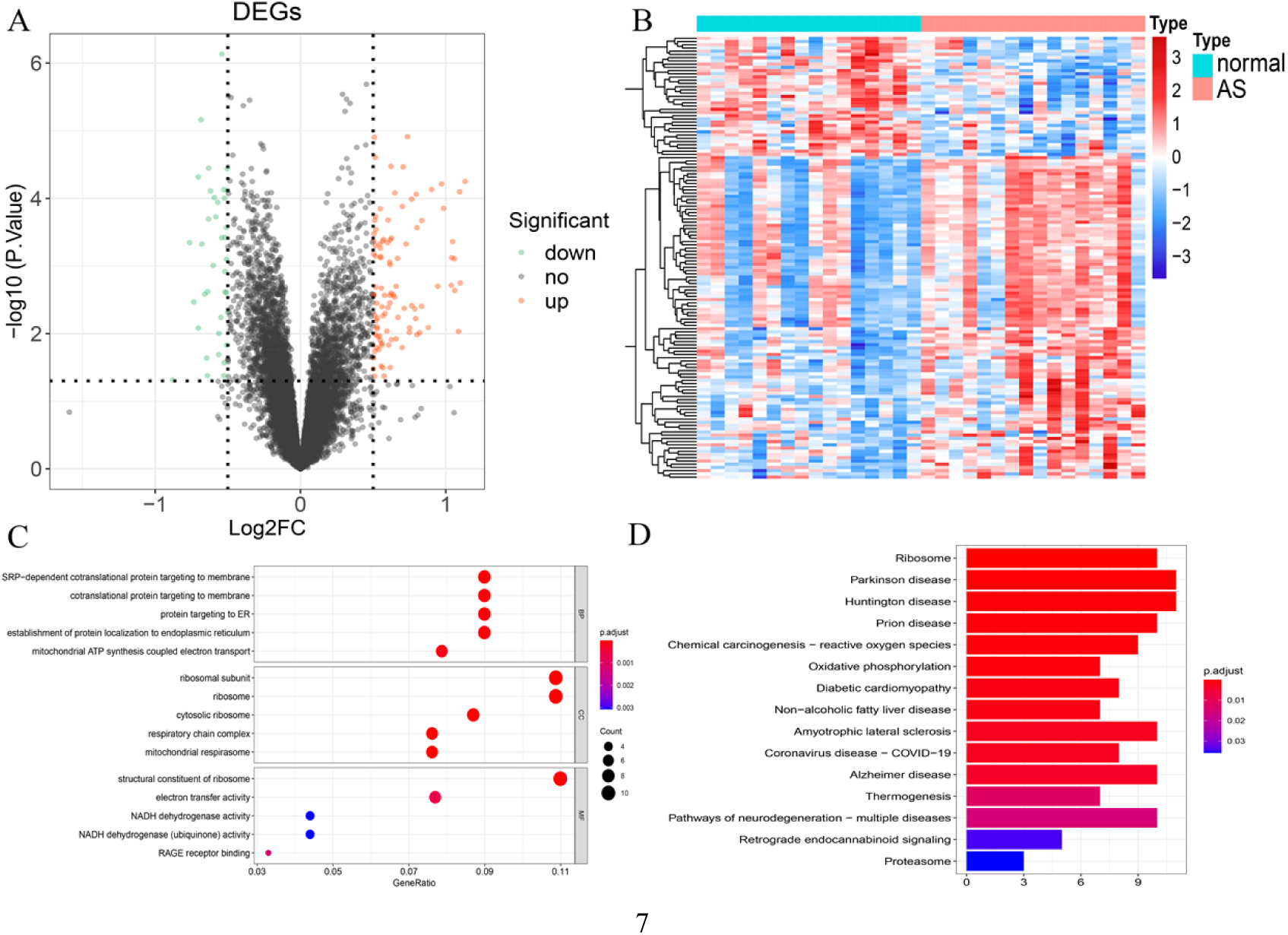
Identification and functional enrichment analysis of DEGs involved in AS. A. The volcano plot of DEGs identified inGSE25101 (n=32, p < 0.05) ; B. The heatmap plot of DEGs identified identified in GSE25101; C. The volcano plot of DEGs identified in GSE25101 (p < 0.05) ; D. Enrichment analysis of GSE25101.

### 8 key common genes were identified in UC and AS

By overlapping those DEGs, 20 common DEGs were found in UC and AS (Fig 3A), including DGKQ, PPBP, GLRX, CKLF, RGS18, S100A8, TMEM158, CLEC4D, C19orf59, CMTM2, S100A12, GNG11, GMFG, LY96, TSTA3, SLPI, S100P, ANKRD22, VAMP5 and HDC, most of which were significantly up-regulated in UC and AS samples (Fig 3B and 3C). To evaluate their performance in classifying UC or AS samples from control ones, we performed ROC curves. As shown in Fig 3D, DGKQ, PPBP, GLRX, S100A8, TMEM158, CLEC4D, CMTM2, S100A12, GNG11, GMFG, LY96, TSTA3, SLPI, S100P, ANKRD22 and VAMP5 may serve as diagnostic markers for UC in GSE87466. As shown in Fig 3E, GMFG, GNG11, CKLF, CLEC4D, CMTM2, VAMP5, S100A8, S100A12 and DGKQ had better performance with AUC greater than 0.8 in distinguishing AS samples in GSE25101. The intersection of those genes, including GMFG, GNG11, CLEC4D, CMTM2, VAMP5, S100A8, S100A12 and DGKQ, were then identified as key common genes in UC and AS (Fig 3F).

**Fig 3.**
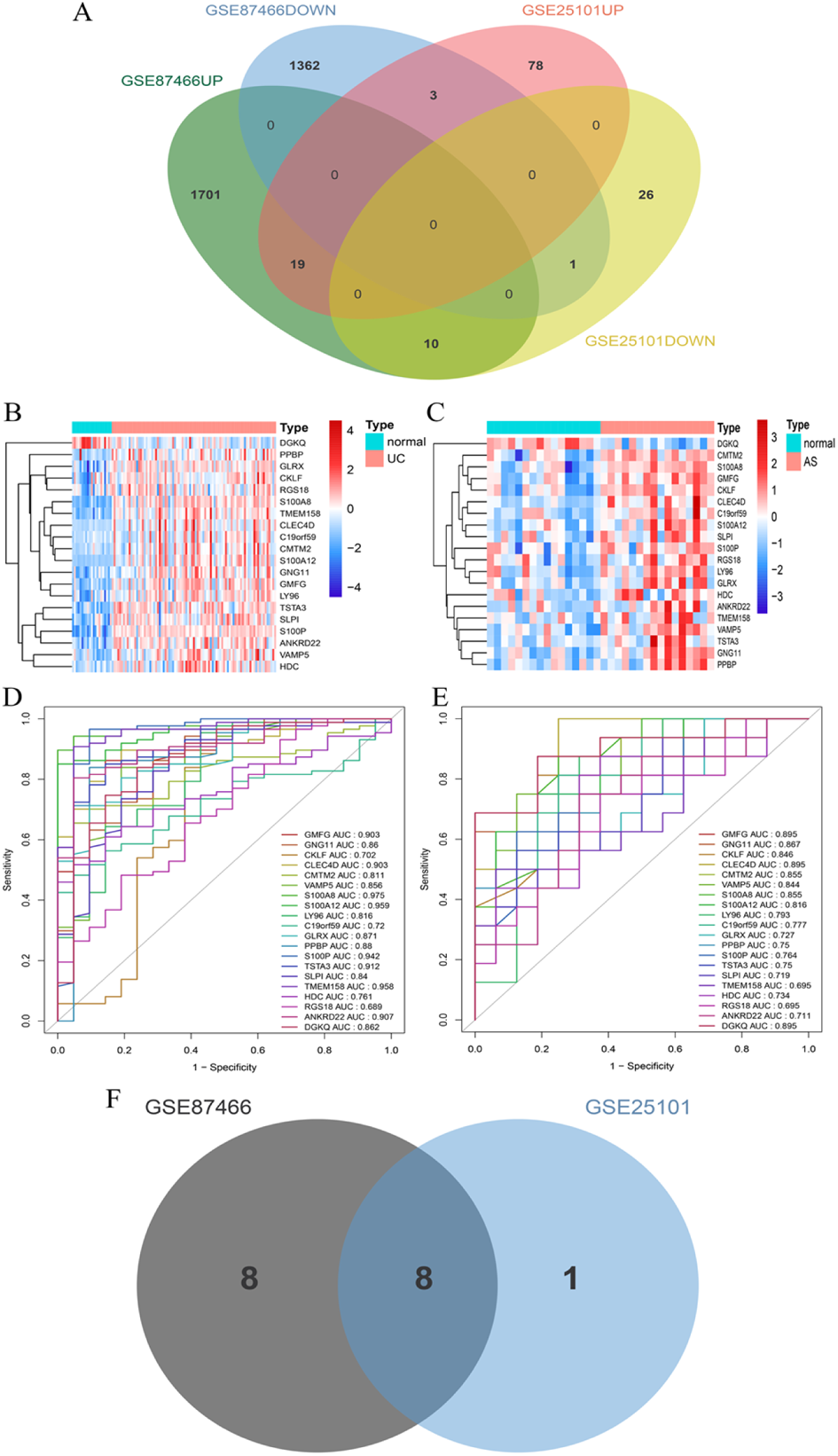
Evaluation and the diagnostic value of common DEGs involved in both UC and AS. A.The Venn diagram displays the interaction of 20 genes within the modules of both UC and AS; B.The heatmap plot of DEGsAU in GSE87466;C. The heatmap plot of DEGsAU in GSE25101; D. ROC curve of 20 DEGsAU in GSE87466 E. ROC curve of 20 DEGsAU in GSE25101; F. The Venn diagram of genes with diagnostic potency in GSE87466 and GSE25101 (n=8).

### Key common genes were regulated by multiple miRNAs and TFs

The miRNAs that target the key common genes were predicted using miRWalk database, and this led to the construction of miRNA-key gene network that composed of 52 nodes and 45 edges (Fig 4A). TFs regulating the expressions of key genes were predicted by Cistrome database, and led to the establishment of a TF-key gene network composed of 167 nodes and 230 edges (Fig 4B). Moreover, the TransmiR database was utilized to identify the relationships between TFs and miRNAs. By integrating miRNA-key gene, TF-key gene and TF-miRNA pairs, a TF-miRNA-key gene regulatory network with 212 nodes and 721 edges was created (Fig 4C). In addition, DGIdb database was utilized to identify potential medicines that target S100A12 and S100A8 for the treatment of UC and AS (rimegepant, eptinezumab, methotrexate, atogepant and ubrogepant) (Fig 4D).

**Fig 4.**
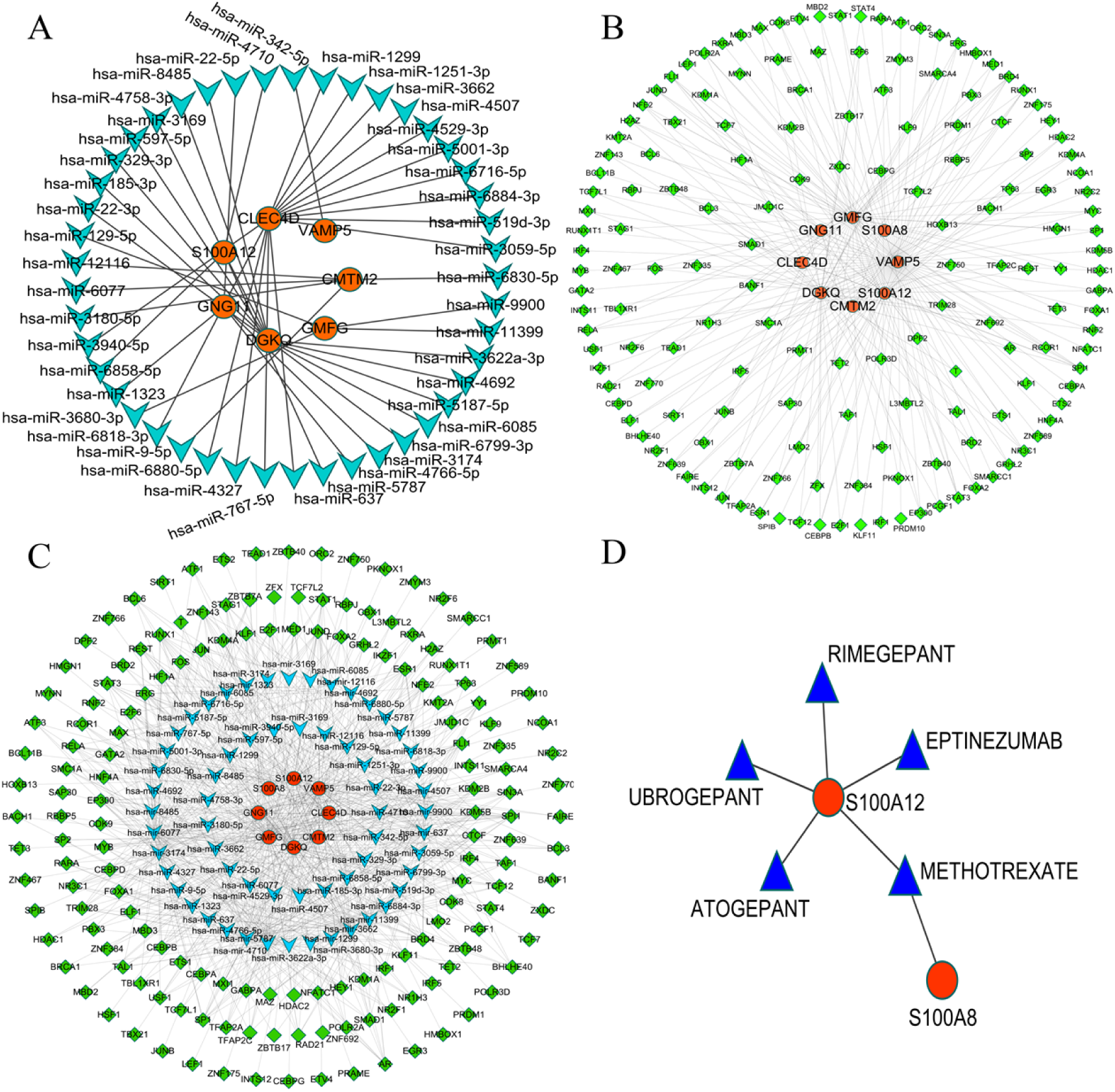
TF, miRNA and potential drugs for key genes involved in both UC and AS. A. miRNA-mRNA co-expression network; B. mRNA-TF co-expression network; C. mRNA-miRNA-TF co-expression network; D. key genes and drugs network.

### 7 common pathways associated with key genes were involved in UC and AS

GSEA analysis was conducted to investigate the potential processes of those key genes in regulating UC and AS. As shown in S7 table and S8 table, the analysis revealed those key genes were linked with various immune related pathways in both AS and UC, comprising antigen processing and presentation, natural killer cell mediated cytotoxicity and T cell receptor signaling pathway, et al. The top 9 enriched KEGG pathways associated with each key gene in AS and UC were displayed in Fig 5 and Fig 6, respectively. Meanwhile, pathways associated with AS and UC were also searched via CTD database (S9 Table and S10 Table). The top 9 pathways related with AS were NOD-like receptor signaling pathway, RIG-`I` -like receptor signaling pathway, JAK/STAT signaling pathway, HTLV-`I` infection, inflammatory bowel disease, Herpes simplex infection, amoebiasis, allograft rejection and cytokine-cytokine receptor interaction (Fig 7A), and the top 9 pathways conferred to UC were TNF signaling pathway, pathways in cancer, chemokine signaling pathway, amoebiasis, AGE-RAGE signaling pathway in diabetic complications, toxoplasmosis, inflammatory bowel disease, tuberculosis and cytokine-cytokine receptor interaction (Fig 7B). Intersecting the results from GSEA and CTD revealed key pathways associated with regulating UC and AS, These pathways include Toll-like receptor signaling pathway, Antigen processing and presentation, allograft rejection, natural killer cell mediated cytotoxicity, pathways in cancer,graft versus host disease and viral myocarditis were identified as key pathways associated with key genes in regulating UC and AS (Fig 7C). In addition, the key genes involved in regulating had high interaction scores with AS related nephritis and UC related systemic lupus erythematosus and hypersensitivity (Fig 8).

**Fig 5.**
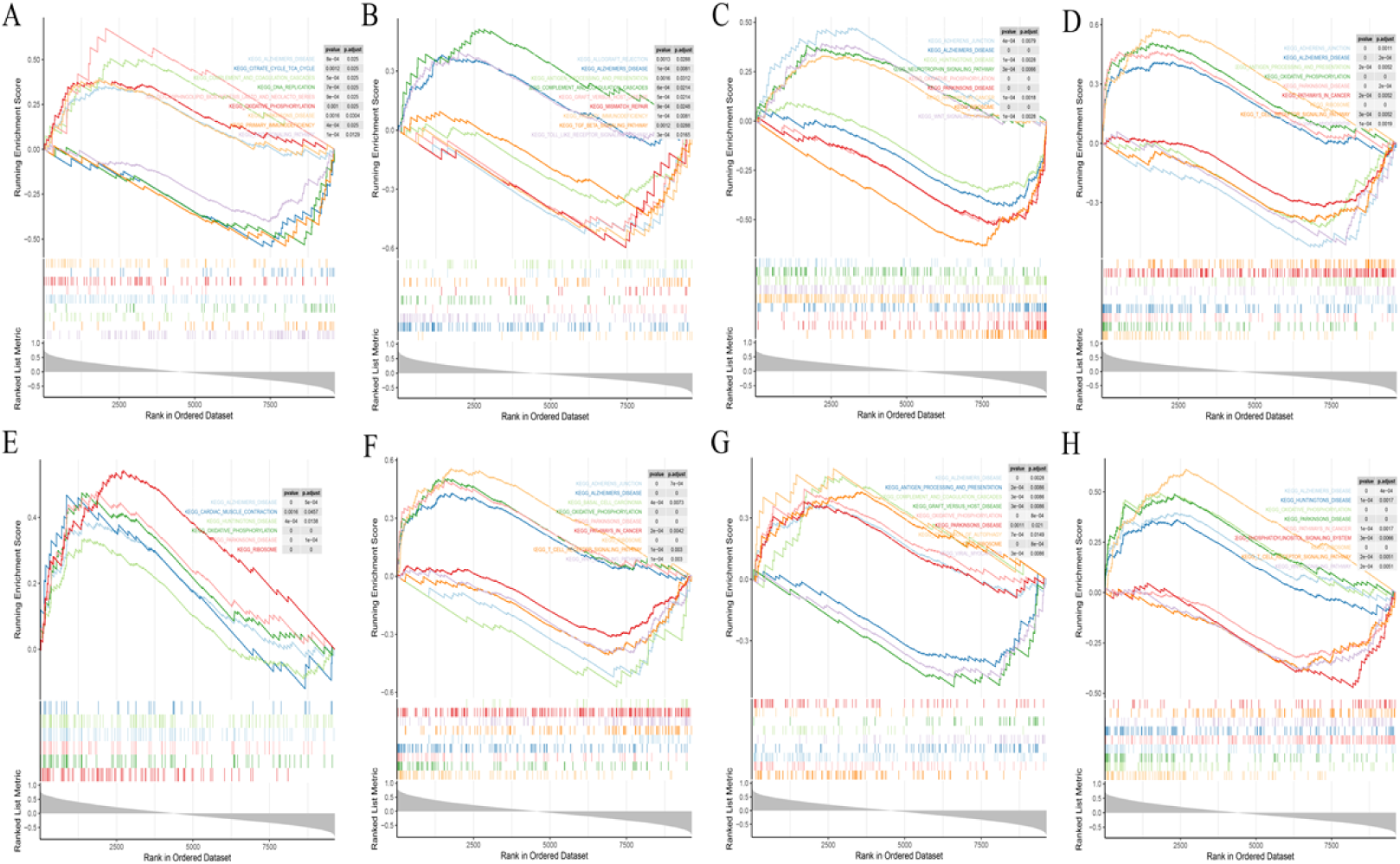
GSEA of the common key genes involved in both UC and AS- GSE25101. A. GSEA of CLEC4D in AS and UC-GSE25101; B. GSEA of CMTM2 in AS and UC-GSE25101; C. GSEA of DGKQ in AS and UC - GSE25101; D. GSEA of GMFG in AS and UC-GSE25101; E. GSEA of GNG11 in AS and UC - GSE25101; F. GSEA of S100A8 in AS and UC-GSE25101; G. GSEA of S100A12 in AS and UC - GSE25101; H. GSEA of VAMP5 in AS andUC-GSE25101.

**Fig 6.**
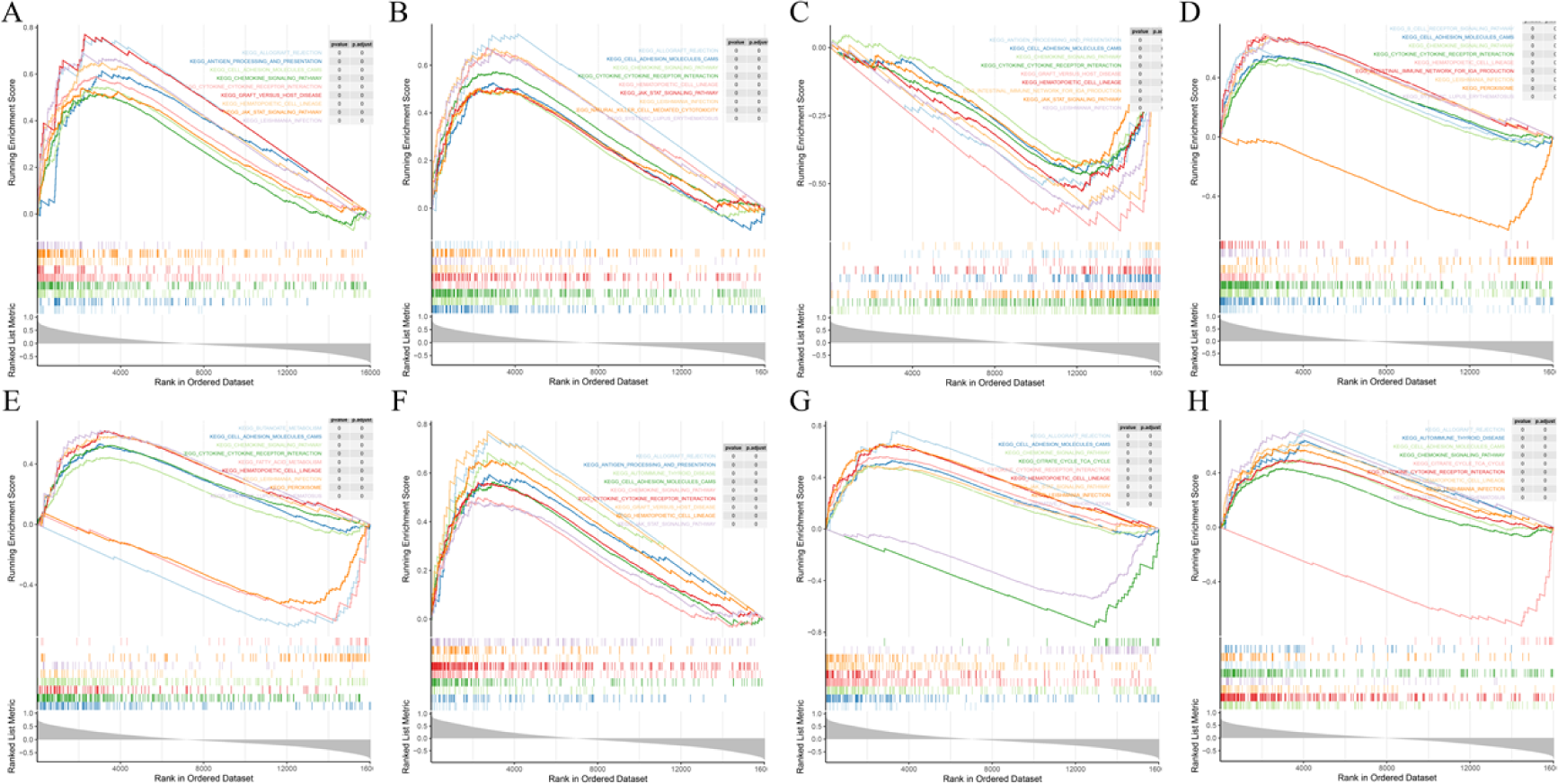
GSEA of the common key genes involved in AS and UC - GSE87466. A. GSEA of CLEC4D in AS and UC - GSE87466; B. GSEA of CMTM2 in AS andUC-GSE87466; C. GSEA of DGKQ in AS and UC - GSE87466; D. GSEA of GMFG in AS andUC-GSE87466 ; E. GSEA of GNG11 in AS and UC - GSE87466; F. GSEA of S100A8 in AS andUC-GSE87466; G. GSEA of S100A12 in AS and UC - GSE87466; H. GSEA of VAMP5 in AS andUC-GSE87466.

**Fig 7.**
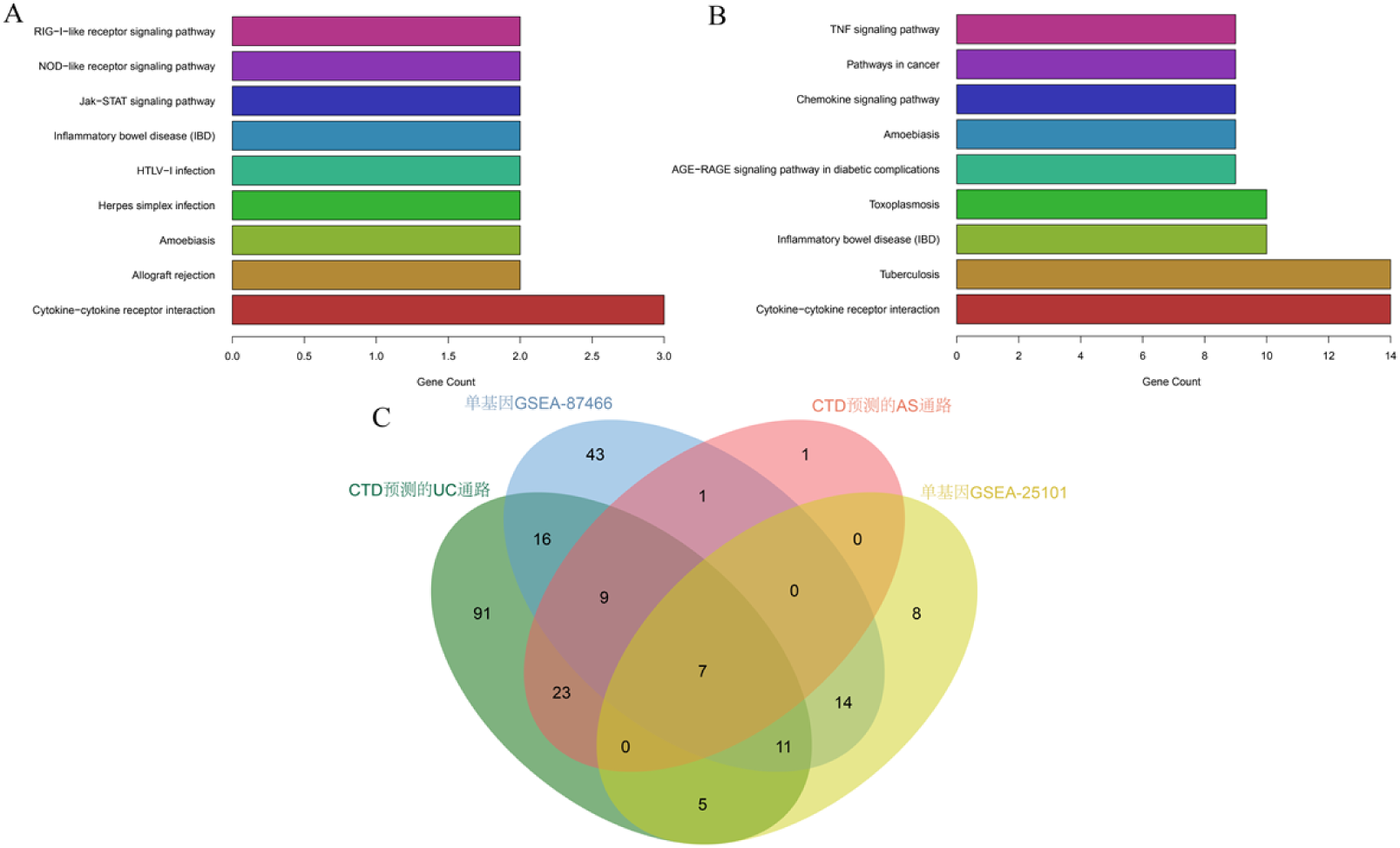
KEGG pathways associated with both AS and UC. A. KEGG path associated with both AS and UC - AS; B. KEGG path associated with both AS and UC - UC; C. The Venn diagram displays the intersection of KEGG pathways in UC and AS.

**Fig 8.**
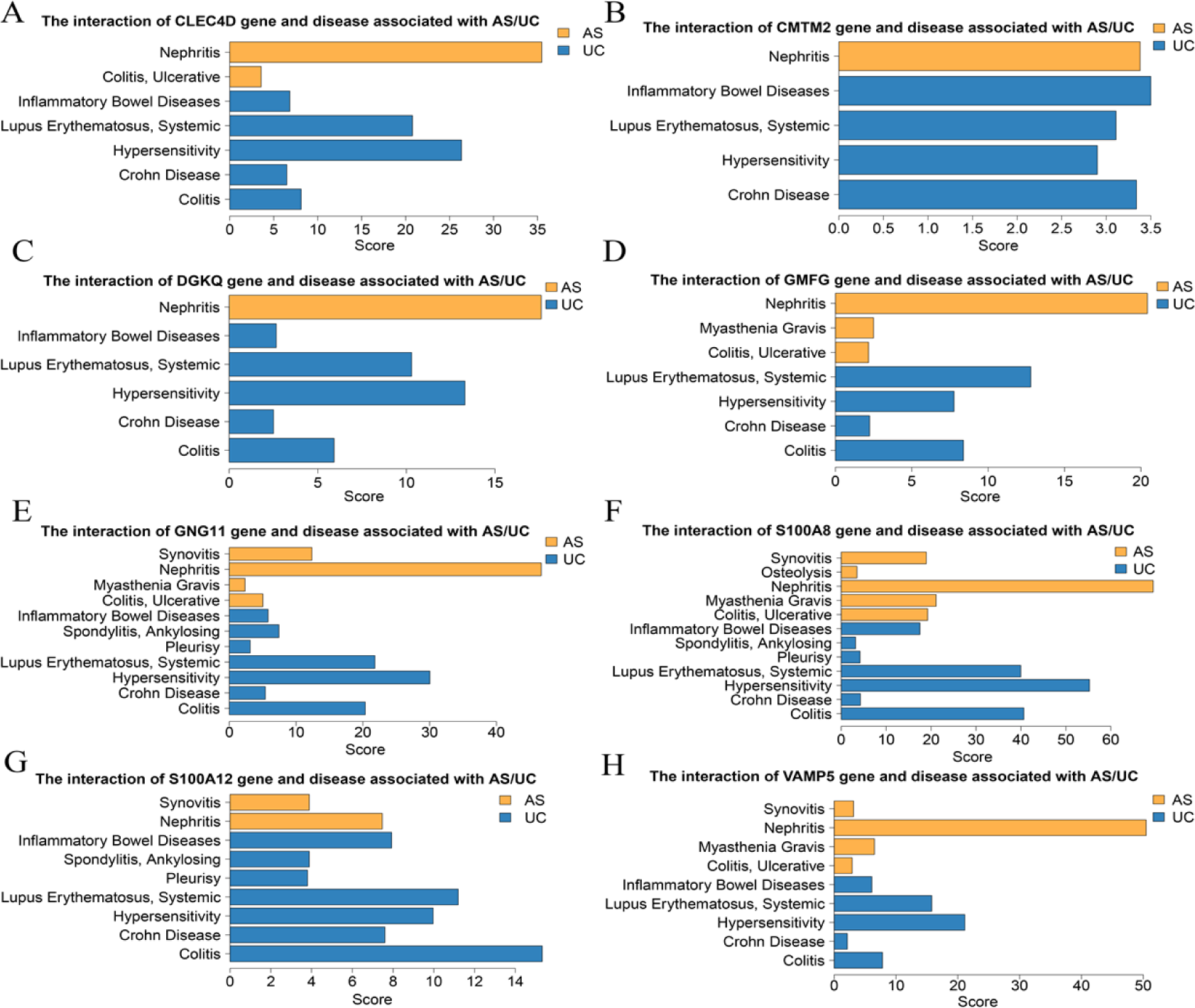
the interaction of key genes and disease associated with AS/UC.

## Discussion

The interrelationship between AS and UC has been widely reported. Combining attacks of the two diseases can result in severe symptoms and poor prognosis, which are frequently misdiagnosed and undertreated[11]. To establish a theoretical foundation for early diagnosis and treatment, we conducted this experiment to investigate the potential genes and pathways being shared in the development of both diseases. Our study identified eight key genes (GMFG, GNG11, CLEC4D, CMTM2, VAMP5, S100A8, S100A12 and DGKQ) involved in AS and UC, and the ROC results indicated that they could be used as potential diagnostic genes for AS and UC. In addition, through intersecting the results from GSEA and CTD, Toll-like receptor signaling pathway, antigen processing and presentation, allograft rejection, viral myocarditis, pathways in cancer, graft versus host disease and natural killer cell mediated cytotoxicity were identified as key pathways associated with key genes in regulating UC and AS.

Glia maturation factor gamma (GMFG) protein is predominantly expressed in immune cells, which has been implicated to play a role in actin reorganization, neutrophil chemotaxis, T cell adherence and so on[12]. A recent study has demonstrated a positive connection between GMF-γ expression and the inflammatory process of UC[13]. The involvement of GMFG in the TLR4 signaling pathway (LPS-induced) was investigated in THP-1 and human primary macrophages[14]. GMFG is revealed to negatively regulate the MAPK, NF-κB, and IFN regulatory factor 3 signaling pathways[15]. knocking down the GMFG gene in macrophages led to elevated levels of mitochondrial reactive oxygen species (mtROS) which was linked to reduced levels of several mitochondrial respiration chain components, including the antioxidant enzymes (SOD1 and SOD2) and the iron-sulfur cluster assembly scaffold protein ISCU[16]. It has been revealed that oxidative stress and the ROS signaling pathway contribute to both AS and UC by enhancing the release of pro-inflammatory cytokines (IL-1β, IL-6 and TNF-α)[17]. Our study shows that GMFG is a common critical gene in both AS and UC, the aberrant linkages between GMFG and the ROS signaling pathway might contribute to the development of UC and AS.

GNG11, a member of the gamma subunit family of G-protein, has been discovered to be expressed in macrophages and dendritic cells. In a gene array study, GNG11 was shown to be associated with psoriatic arthritis (PsA) individuals compared to healthy controls[18]. It appears to respond to oxidative stress by inducing its gene expression. Moreover, GNG11 plays a role in transmitting external signals to internal effectors, and its over-expression has been firmly linked to the onset of cellular senescence through activating the extracellular signal-regulated kinase (ERK)1/2 pathway[19]. ERKs signaling pathway is one of the best-characterized MAPK signaling pathways. Through phosphorylating transcription factors like c-Fos and c-Jun, activated ERK can directly influence the cell division and proliferation[19]. Moreover, it has been demonstrated that ERK1/2 activation promotes osteoclast formation and bone resorption, which may aid in the development of spinal abnormalities and bone erosions in AS[20, 21]. The aetiology of UC is closely tied to the traditional mechanism for ERK pathway. After being activated, ERK may regulates its downstream targets, such as NF-κB and Bcl-2, which can influence the inflammatory response and cell death in the development of UC[22]. According to the above statements, our research indicates that GNG11 is a crucial gene shared by both AS and UC, and it may contribute the co-pathogenesis of AS and UC by influencing the concentrations of associated transcription factors via ERK1/2 pathway.

CLEC4D (also known as Dectin-3/ MCL) which is a member of Myeloid C-type lectin receptors (CLRs), could recognize microbial components and subsequently activate intracellular signaling pathways while modulating the immune responses[23, 24]. CLRs exhibit diverse functions, such as inducing endocytic, phagocytic, antimicrobial, pro-inflammatory and anti-inflammatory responses, and these functions are dependent on the particular signaling motifs found within the cytoplasmic domains of the CLRs. The CLRs transmit signals via the spleen tyrosine kinase (Syk), which then combines with the CARD9/ B-Cell CLL/ BCL-10/ MALT1 complex. This results in the release of pro-inflammatory cytokines and activation of adaptive T-cell immunity[23]. CLEC4D predominantly expresses in macrophages, neutrophils in the peripheral circulation, classical monocytes, and certain subsets of dendritic cells (DCs). It plays a role in endocytosis and engages to mycobacteria by recognizing TDM on the cell walls[25]. Binding to the TDM can upregulate the expression of Mincle, leading to an enhanced cellular response. The downstream signaling is mediated by CARD9/BCL10/MALT1 and Syk kinase pathways. These pathways induce numerous intracellular responses, including the activation of NF-κβ, phagocytosis, and the release of inflammatory cytokines[26, 27]. Moreover, the interaction of CLEC4D with TDM promotes the maturation of DCs and the priming of T cells within the body[28]. A higher level of CLEC4D mRNA was revealed in patients with AS using whole blood transcriptional profiling[29]. Furthermore, it was found that CLEC4D-deficient mice were more vulnerable to colitis than wild-type mice (induced by DSS)[30]. CLEC4D gene polymorphisms have been connected with TB prevalence, making the CLR a key component of anti-mycobacterial defense[31]. The association between Syk and the initiation of inflammation in osteoarthritis (OA) has been revealed, resulting in the secretion of inflammatory factors such as IL-1β that hasten the degradation of cartilage and bone tissue. The severity of the inflammatory response can be decreased by using inhibitors of Syk[32, 33]. As a clinical focus of UC, Syk participates in the initial stages of B-cell receptor and Fc receptor signaling[34]. In a mouse model of colitis induced by acetic acid, the selective Syk inhibitor (fostamatinib) resulted in a reduction of mucosal damage[35]. In our study, CLEC4D was found to be elevated in UC and AS by overlapping the DEGs. Although there are limited studies on the association of CLEC4D gene with AS and UC, the aforementioned researches allow us to speculate on the potential mechanisms of CLEC4D in AS and UC. By stimulating the Syk kinase/ CARD9/ BCL10/ MALT1 pathway and Mincle transcription, which in turn activates NF-κβ and pro-inflammatory cytokines (IL-1β, IL-6 and TNF-α), CLEC4D may accelerate the development of AS and UC.

Chemokine-like factor (CKLF) superfamily (CKLFSF), also known as CMTM, is a critical component in both pathological and physiological processes. In recent years, CMTM has gained the interests of researchers due to its potential effects on a wide range of disorders. CMTM2, a sub-types of CMTM, was primarily found in bone marrow and CD4+T cells[36]. Multiple studies have indicated that CMTM2 might play a role in suppressing HIV-1 LTR activity by blocking the CREB and AP-1 pathways[37]. According to a study in Chinese cohort, individuals with HBV-related liver disorder exhibited notably lower expression levels of CMTM2 in their serum compared to healthy subjects. In addition, it has been demonstrated that serum CMTM2 correlates with the DNA load of HBV and may be utilized to distinguish between HBV patients and healthy individuals[38]. Comprehensive histologic and microarray analysis of OA patients revealed reduced CMTM2 expression in OA patients compared to healthy controls[39]. Furthermore, the utilization of whole-genome microarray revealed that AS patients exhibited over-expression of CMTM2 in their peripheral blood[29]. Our study found that CTMT2 was up-regulated in both UC and AS, based on the overlap of DEGs. Despite the limited studies on the association between the CTMT2 gene with AS and UC, our findings provide insights into the potential pathways of CTMT2 in these two conditions.

VAMP5 is an important member of the Soluble N-Ethylmaleimide Sensitive Factor (SNARE) protein family[40], which plays a role in vesicle docking and fusion by interacting with other SNAREs. Vesicle-mediated protein transportation is the primary means of transporting proteins along the secretory and endocytic pathways, and which regulates the secretion of proteins and neurotransmitter release[41]. This process is essential for the proper functioning of cells, as it allows proteins to be delivered to the correct locations in the cell, and regulates the release of neurotransmitters. VAMP5 is an important gene in facilitating intracellular transport functions such as exocytosis, internal recycling and endocytosis. Mice lacking VAMP5 had a low birth rate and respiratory systems defects, indicating its functional role in developing respiratory system[42]. The elevated expression of the genes in tuberculosis patients indicates their crucial roles in the disease’s immunological mechanisms. researches have demonstrated that the levels of VAMP5 are relatively higher in CD14+ and CD16+ monocytes compared to other cell types. This aligns with the role of monocytes in tuberculosis infection, as they participate in vesicle transportation, which require VAMP5 for this process[42, 43]. Furthermore, the elimination of tuberculosis could involve the complement activation pathway, and the predicted role of the discovered protein in tuberculosis is closely linked to its sequence-structure-function relationship[44]. Multiple genetic variation studies suggested that the VAMP5 gene conferred the susceptibility of Hirschsprung disease (HSCR), demonstrating the involvement of VAMP5 in the physiological processes of the digestive system[45]. Our study identified VAMP5 as a common key gene in UC and AS. Given the paucity of data on this topic, further researches are required to confirm the underlying link between the VAMP5 gene and AS with UC illness.

Diacylglycerol kinases (DGKs) are a class of enzymes that catalyse the phosphorylation of diacylglycerol (DAG) to form phosphatidic acid (PA) which is a vital intermediate of lipid metabolism and signaling pathways within cells[46]. DGKQ, a critical member of DGKs, consists of three cysteine-rich domains (CRDs), a proline/glycine-rich region, and a pleckstrin homology (PH) domain that overlaps with a Ras-binding domain[47]. Previous studies have showed that DGKQ might be linked to several autoimmune diseases, including systemic lupus erythematosus (SLE), type 1 diabetes (T1D), RA, idiopathic inflammatory myopathies (IIM), primary biliary cirrhosis (PBC), Sjögren’s syndrome (SS) and so on[48–50]. The high-risk SNP alleles were found to be linked with increased expression of DGKQ in fibroblasts, lymphocytes and lung. The DGKQ protein serves as a critical mediator for cell signal transduction and is expressed at higher levels in the vasculature of systemic sclerosis patients with pulmonary involvement[51]. This protein can indirectly stimulate the epidermal growth factor receptor (EGFR) pathway[52], which is crucial for regulating cell proliferation and migration. Our study identified DGKQ as a crucial gene shared by UC and AS. To confirm this result, additional research must be conducted.

The S100 family proteins comprises 25 recognized members that exhibit a considerable level of structural and sequence similarity. Interacting with signal transducers and various receptors, they modulate the processes that involve in inflammation, energy consumption, calcium homeostasis, apoptosis, cell cytoskeleton, microbial resistance, cell differentiation and proliferation[53]. In our research, both S100A12 and S100A8 were identified as common key genes for AS and UC. S100A8, a type of Ca^2+^-binding protein, is highly expressed in a variety of infectious and inflammatory disorders including RA, IBD, pseudomonas aeruginosa keratitis psoriasis and acute anterior uveitis (AAU) [54–56]. In addition, S100A8 (also known as MRP8) primarily engaged to S100A9 (also known as MRP14) and formed a heterodimer[56, 57]. By combining to the receptor of advanced glycation endproducts (RAGE) and TLR4, extracellular S100A8/S100A9 is also hypothesised to affect a number of processes in the leukocyte recruitment. As a result, the transcription factor NF-kB is activated, which leads to the release of pro-inflammatory cytokines (IL-1β, IL-6 and TNFα)[58]. Moreover, the activation of S100A8/S100A9 in monocytes may promote ROS to enhance the production of pro-inflammatory cytokines and the expression of NLRP3 inflammasome[59, 60]. An increased level of S100A8 has been revealed in the patients with arthritis and IBD, indicating that it could play a role in AS and UC by affecting the aforementioned pathways and cytokines.

S100A12, which is an alarm signal that specifically targets granulocytes, binds to RAGE and TLR4[61]. According to reports, RAGE-dependent activation of NF-κB has been shown to result in the release of pro-inflammatory cytokines, ultimately leading to the recruitment of monocytes[62]. S100A12 is involved in the recruitment of inflammatory cells in mouse models and has been reported to be over-expressed in inflamed tissues of patients with a variety of conditions, including IBD, PsA, JIA and RA[63–65]. It is considered as a reliable biomarker for IBD and systemic-onset JIA[66]. Both JIA and AS are thought to be autoimmune illnesses in which the immune system mistakenly targets the body’s own tissues, resulting in joint inflammation and damage. This implies S100A12 may play a critical role in the development of IBD and AS.

Therapeutic agents that serve as functional inhibitors of receptors and ligands have been widely employed to managing diverse conditions, including immunological disorders. Studies have shown that by binding to S100A12, anti-allergic medication can effectively block downstream RAGE signaling and NF-κB activation[67]. According to research conducted on mice models, the use of inhibitors and antibodies to limit the activity of S100A8/A9 can improve pathological conditions[54]. Certainquinoline-3-carboxamides, that are widely studied for the treatment of human autoimmune and inflammatory illnesses, interact with S100A9 and the S100A8/A9 complex, thereby reducing their interaction with TLR4 or RAGE and TNF-production in vivo[68]. A recent study indicates that Blocking S100A8/A9 reduces inflammatory processes in mice arthritis models[69]. It has been suggested that targeting S100A8 could be an effective strategy to address the chronic inflammation caused by obesity. By DGIdb database, we found that rimegepant, eptinezumab, methotrexate, atogepant and ubrogepant may serve as potential drugs for treating UC and AS via targeting S100A12 and S100A8. Given the possible significance of S100L8 and S100L12 genes to AS and UC, it is reasonable to assume that these medications could serve as research topics for disease treatment in the future.

In the present study, GSEA was performed to identify the pathways that the keys genes exhibited significant enrichment. GMFG, CLEC4D, S100A8 and S100A12 exhibited high levels of enrichment in various processes related to the activation and function of neutrophils, including neutrophil activation involved in immune response, neutrophil mediated immunity, neutrophil degranulation, and neutrophil activation pathways (shown in Table S5 and S6). Neutrophils have long been recognized as important players in the immune system, both in terms of innate and adaptive immunity. Recent research has revealed that neutrophils can exhibit pronounced phenotypic and functional abnormalities in a variety of systemic autoimmune disorders. These abnormalities can lead to inappropriate immune responses and organ damages in these diseases. Both GMFG and S100A8 were showed to be connected with the Toll-like receptor pathway which has been identified as a crucial role in numerous autoimmune diseases. In our study, Toll-like receptor pathway was also identified as a common process of AS and UC. In addition, we revealed natural killer cell mediated cytotoxicity, antigen processing and presentation, allograft rejection, pathways in cancer, viral myocarditis and graft versus host disease as key pathways associated with aforementioned genes in regulating UC and AS. These pathways discovered above have provided us with new ideas for future research.

## Materials and Methods

### Data source

In this study, gene expression profiles of 21 control and 87 UC samples in GSE87466 dataset based on GPL13158 platform, and 16 control and 16 AS samples in GSE25101 dataset based on GPL6947 platform were acquired from GEO database (https://www.ncbi.nlm.nih.gov/geo/).

### Identification and functional enrichment analysis of DEGs

Limma R package was employed to detect differentially expressed genes (DEGs) between control and UC subjects in GSE87466 and DEGs between control and AS samples in GSE25101 with |log_2_FC| >0.5 and adjusted p-value <0.05. The findings were displayed in volcano plot and heatmap generated by ggplot and pheatmap R package, respectively. ClusterProfiler R package was utilized for enrichment analysis of GO and KEGG pathway in DEGs (Figure 1 and Figure 2). In the GO analysis, Biological process (BP), cellular component (CC) and molecular function (MF) were included. And a p-value <0.05 was regarded as significantly enriched. By overlapping DEGs in GSE87466 and GSE25101, common DEGs involved in both UC and AS were identified and used for the downstream analysis (Figure 3).

### Evaluation of the diagnostic value of common DEGs

ROC curves were conducted to evaluate the performance of common DEGs in classifying AS or UC samples from control ones. Common DEGs with AUC value greater than 0.8 were defined as potential diagnostic biomarkers for AS or UC(Figure 3D and Figure 3E). Then we overlapped the diagnostic biomarkers for AS and UC and identified them as key genes connecting AS and UC (Figure 3F). The miRWalk (http://mirwalk.umm.uni-heidelberg.de/), Cistrome (http://cistrome.org/) and TransmiR (http://www.cuilab.cn/transmir) to search for miRNA-key gene, TF-key gene and TF-miRNA relationships and to construct a TF-miRNA-key gene regulatory network by Cytoscape software (Figure 4A, Figure 4B and Figure 4C). In addition, potential drugs for key genes were predicted by DGIdb database (Figure 4D; https://www.dgidb.org/).

### Identification of key genes-related mechanisms underlying in UC and AS

According to the correlations with each key gene, genes in GSE87466 and GSE25101 were ranked and GSEA were performed to detect the pathways that those genes were significantly enriched into (Figure 5 and Figure 6). KEGG pathway gene sets were obtained from MSigDB database (Figure 7, https://www.gsea-msigdb.org/gsea/msigd b) and were used as reference gene sets. In addition, AS and UC related pathways were also downloaded from the CTD database (http://ctdbase.org/) using the key words “ulcerative colitis” and “ankylosing spondylitis”. By intersecting pathways identified by GSEA and CTD, key gene-related pathways involved in UC and AS were identified. Meanwhile, via CTD database, we also investigated the interaction of each key gene and diseases associated with UC or AS.

## Conclusions

For the first time, our study highlighted 8 common key genes (GMFG, GNG11, CLEC4D, CMTM2, VAMP5, S100A8, S100A12 and DGKQ) and 7 common pathways (Toll-like receptor signaling pathway, antigen processing and presentation, allograft rejection, viral myocarditis, pathways in cancer, graft versus host disease and natural killer cell mediated cytotoxicity) shared in UC and AS. Understanding the roles of these genes and pathways in shaping autoimmunity offers great potential for the future therapeutic development in AS and UC intervention.

## Acknowledgements

This work of Liping Du and Xuemin Jin was supported by National Natural Science Foundation of China [819707922 and 82171040], Medical Science and Technology Project of Health Commission of Henan Province [YXKC2020026], Key Scientific Research Projects of Henan Province Colleges and Universities [23A320067]. The work of Fuzhen Li was supported by Major Program of Medical Scientific and Technological of Henan Province (XXX), Medical Scientific and Technological Project of Henan Province (SBGJ2020003031) and National Natural Science Foundation Project (82101108).

## Authors’ contributions

LL, FL, XJ, LD and PY designed the study. LL, FL, KX and PZ downloaded the raw data. LL, FL and HZ conducted the analysis. LL drew up the initial draft. XJ, PY and LD reviewed the manuscript. All authors read and approved the final manuscript.

